# Investigation of Regulation and Binding Patterns of the Human Cathelicidin Peptide LL-37 in Complexation with Nucleic Acids, and its Impact on Neutrophil Extracellular Traps

**DOI:** 10.64898/2026.02.09.704888

**Authors:** Claudia Zielke, Behzad Rad, Josefine E. Nielsen, Jiaxin Li, Sopida Pimcharoen, Manasi Sawant, Jennifer S. Lin, Hawa R. Thiam, Annelise E. Barron

## Abstract

The human cathelicidin host defense peptide LL-37 forms complexes with nucleic acids that can have either beneficial or detrimental health effects. We suggest that these differential impacts are directly connected to dsDNA binding by LL-37 and to complex formation between protomers. Here, we show using phage λ DNA that LL-37 binds non-specifically to dsDNA, condensing it, followed by complex formation between LL-37 peptides. We find that complex formation is concentration-dependent, with low LL-37 amounts yielding loosely aggregated DNA structures, while higher LL-37 concentrations lead to well-defined, disc-like structures of about 150 nm in diameter. The condensation of the nucleic acids, which causes a loss of the characteristic B-DNA features, results from interactions of the phosphodiester backbone with protonated amino acid side chains of the peptide at physiological pH, predominantly in A-T rich sequences of the nucleic acid. However, in our studies, electrostatic interactions did not appear to be the driving force for complexation, but rather we found the α-helical structure of the peptide with its amphipathic and hydrophobic surfaces to be essential. Further, we show that LL-37 also interacts with nucleic acids from neutrophil extracellular traps (NETs) in a concentration-dependent way, causing a reduction in NET aggregate area, which may offer new biophysical insights into diseases such as systemic lupus erythematosus (SLE), which involve slower-than-normal NET clearance. Our results indicate the key importance of LL-37 expression levels for regulation of the innate immune system for optimal human health, since the relative amounts of expressed LL-37 present to interact with extracellular DNA will determine the extent to which the DNA can be condensed, which in turn will affect the ability of the body to clear the NETs before they can cause inflammatory conditions.

## 1. Introduction

The innate immune system is crucial as the first line of defense against invading pathogens such as bacteria and viruses that can infect the human body. Different classes of antimicrobial peptides (AMPs) play a special role in the innate immune system by protecting the body against a broad range of microbial infections. The AMP LL-37 is the sole human cathelicidin and has been shown to have antiviral, antifungal, and antibacterial properties.^1,2^ LL-37 is utilized by neutrophils and other immune cells to target and kill infected cells^3,4^, and is necessary for the killing of pathogens and infected host cells.^5^ However, LL-37 also has been closely connected previously with inflammation and autoimmune diseases, as we will discuss below^1^.

The expression of the LL-37 peptide in the human body is triggered by physical wounding, infection, and endoplasmic reticulum stress response, and can be modulated by Vitamin D3 and butyrate.^6,7^ Subsequent upregulation of LL-37 has been suggested to play an important role in human innate immunity, e.g. through reduction of severe COVID-19 pathology after infection^8,9^, hindrance of *P. aeruginosa* biofilm maturation^10^, an ability to drive angiogenesis, and modulation of inflammatory responses.^11^

LL-37 consists of 37 amino acids (net charge of +6) in an α-helical structure, which enables the peptide to bind to other biomolecules in solution (*e.g.*, amyloid-β^12,13^, islet amyloid polypeptide^14^, α-synuclein ^15^, lipid bilayers/phospholipid membranes^16^) and to protect itself from proteolytic degradation. Additionally, the abovementioned net positive charge makes it a natural binding partner for negatively charged molecules such as different types of nucleic acids (DNA, RNA). Complexes formed between LL-37 and nucleic acids have been widely discussed in the literature for their myriad seemingly contradictory impacts on the human body, with both beneficial and detrimental health effects.^17^

In several studies, these LL-37/DNA complexes were shown to be immunostimulatory with significant beneficial impact on the human innate immune system (as seen in four prior studies^18–21^). For example, LL-37 can target the extracellular DNA of mammalian cells, protecting the complexed nucleic acid from serum nuclease degradation^18^ and is important for fundamental processes such as cell signaling, coagulation, and innate immunity.^22^ Zhang *et al.* then described mechanisms of LL-37 functioning as a cargo delivery vehicle for dsDNA through membrane perturbation.^23^ LL-37/DNA complexes were also shown to enhance proliferation and activation of natural killer (NK) cells, subsequently stimulating the innate immune system.^24^ In the latter study, the combination of bound DNA subunits with LL-37 peptides was suggested to control ovarian tumors through the activation of innate immunity.^24^ A recent study found a significant correlation between the levels of the DNA-LL-37 complex and LL-37 within dental plaque (biofilm) specimens^25^. In vitro testing showed these complexes stimulated Toll-like receptor 9 (TLR9). Besides nuclear DNA, LL-37 has been shown to form complexes with a variety of other nucleic acids including RNA^26^ and mitochondrial DNA^27^, and to trigger toll-like receptors TLR3^28,29^, TLR7^26^, TLR8^26^ and TLR9^30,31^, which all play key roles in the innate immune system. In contrast to the above-mentioned beneficial health effects, other studies showed that LL-37/DNA complexes appear to have some involvement in autoimmune diseases such as psoriasis^32,33^, systemic lupus erythematosus^1,34,35^ and atherosclerosis^36^, primarily through the detection of LL-37 peptide within aggregates and complexes associated with disease.

LL-37 can also be found in neutrophil extracellular traps (NETs), which are part of an innate immune response in which genomic DNA is expelled from activated neutrophils to release a web-like structure enabling the NETs to engulf and kill pathogens^37,38^. NETs consist of a DNA/histone backbone associated with cytoplasmic and granule proteins and LL-37.^39,40^ After NET release, NETs need to be cleared by (*e.g.*) DNase I or by macrophages to prevent detrimental NET-related effects such as thrombosis^37,41,42^ or deleterious and inflammatory TLR9 activation. Although a direct link between extracellular LL-37 and NETs has been previously established^43,44^, investigation of the importance and involvement of LL-37 in the biophysics of NET clearance is lacking.

LL-37 dysregulation, *i.e.*, the under- or overexpression of LL-37 in the human body—which is often discussed as being relevant to autoimmune disease—has not been previously considered when talking about binding mechanisms of the complexes. Notably, experimental conditions and the relative molar ratios of peptide to nucleic acids when forming the complexes vary greatly in the literature. These conditions (relative molar ratios of LL-37 to DNA, which would be affected by levels of LL-37 expression, i.e., *CAMP* gene expression) are hypothesized here to influence the biophysical characteristics of LL-37/DNA complexes, and hence their physiological effects. To the best of our knowledge, only a few studies have been published with a structural investigative approach towards the functionality or mechanism of action of LL-37/DNA complexes. Examples seen in literature for LL-37/DNA complex structures depict nucleic acid strands loosely arranged around one LL-37 node to form extended “flower-like” structures^32^, cationic protofibril LL-37 assemblies interacting with anionic DNA to form a square columnar lattice so as to play a role in TLR9 activation^30,31,45^, and spherical nanoparticles of about 120−190 nm in diameter.^46^ A recent review summarized further the powerful immunomodulatory effects of LL-37/DNA complexes in regards to the innate immune system, and discussed the importance of evaluating their molecular structure to be able to draw conclusions on beneficial or detrimental effects on human health.^17^ To yield insights into the interactions of LL-37 with dsDNA, we investigated complexes formed between LL-37 and phage λ DNA at different (w/w) ratios. Phage λ DNA was chosen due to its defined size (48.5 kilobase pairs), reproducibility, and easy handling. We used TIRF microscopy to investigate single dsDNA molecules upon binding LL-37 to understand the condensation patterns of dsDNA under the influence of LL-37. In addition, we use ensemble binding to determine the stoichiometry between LL-37 and dsDNA. Further, we systematically formed LL-37/dsDNA complexes at specific ratios to investigate structures with lower and higher LL-37 concentrations to simulate under- and overexpression of LL-37 in a system. These ratios were then used to analyze the complexes with atomic force microscopy, asymmetrical flow field-flow fractionation with multiple detection and small angle X-ray scattering. Finally, we investigated the impact of LL-37 on NETs; concentration-dependent compaction of the nucleic acids in the NETs may give further insights into the importance of LL-37 regulation of the human body. Altogether, our results indicate LL-37 binds to dsDNA in a strongly concentration-dependent manner, mediating further complex formation and condensation. We believe that the detailed molecular biophysics of LL-37/DNA complex formation regulates the innate immune response and is critical for the physiological effects of LL-37 on human health.

## 2. Results

In order to draw conclusions on the impact of the possible underexpression of LL-37 on DNA complexation within the human body as part of natural innate immune responses, we investigated effects of the binding of LL-37 to λ DNA using different, complementary techniques. Further, the concentration-dependent impact of LL-37 on NETs was studied as well.

### LL-37 condenses λ DNA under a constant flow

Biotinylated λ DNA was attached to a glass-slide enclosed within a flow cell and was subjected to a constant flow of solution consisting of either PBS buffer alone or LL-37 dissolved in PBS (see Materials and Methods). Under an applied flow of PBS only, λ DNA stretches out, as can be seen in **Figure 1A**, where two molecules are stretched out freely next to each other. Under the influence of LL-37 binding, λ DNA strands first interact with each other (**Figure 1B**) followed by condensation of the strands (**Figure 1C**), resulting in a dense structure (**Figure 1D**). This condensation under the given flow conditions was observed to occur within less than 5 seconds. It is evident that the LL-37 peptide has the ability to condense λ DNA to form dense structures, even on a single DNA molecule level.

**Figure 1.**
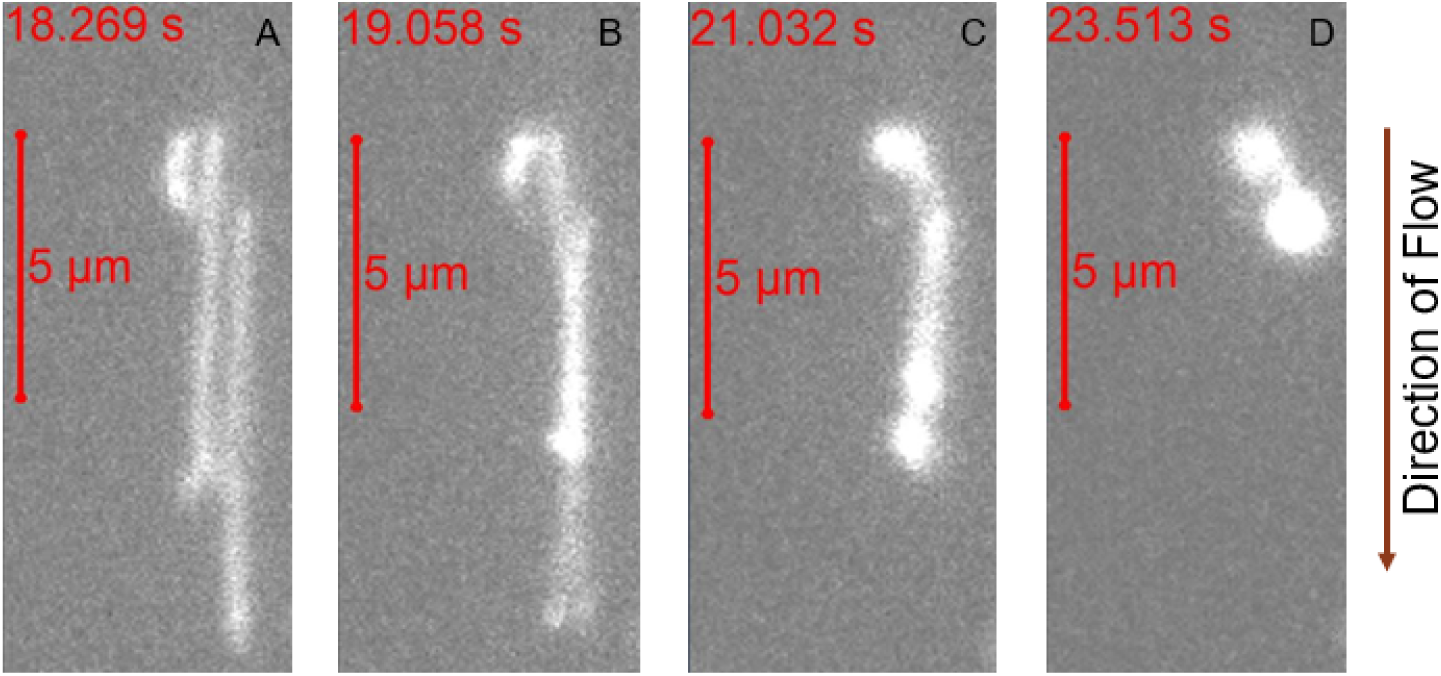
Fluorescently labeled λ-DNA was biotinylated and selectively attached to a glass slide in a flow channel and imaged using high resolution fluorescence microscopy. λ-DNA strands are stretched out under the influence of applied flow and condense to a dense structure under the influence of 1 nM LL-37.

To obtain more data points from this experiment, we used “DNA curtains”, where many DNA strands are attached to a mobile supported lipid bilayer, then arranged under flow at an etched barrier, creating a curtain like set-up.^47^ The same condensation pattern as seen in **Figure 1** for single molecules was observed for λ DNA curtains (**Figure 2**). Under constant flow, DNA stretched out (**Figure 2A**) and interacted with neighboring nucleic acid strands after first contact with LL-37 (**Figure 2B**) before condensing into dense structures (**Figure 2C**). Condensation of strands was analyzed by plotting the length of λ DNA over either time since LL-37 contact, amount of flow running through the chip or LL-37 concentration the nucleic acids have been in contact with at that point of time (**Fig 2D**). Using a Boltzmann Sigmoid Fit according to Equation (1), it was calculated that the nucleic acid is condensed in 3.83 ± 0.08 μm/s before full condensation.

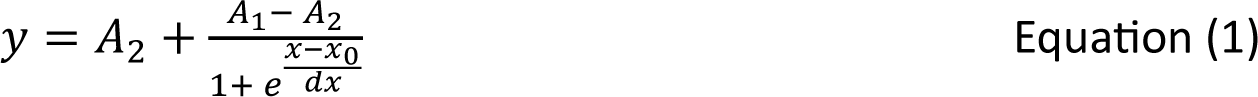

with *dx* being the condensation constant.

**Figure 2.**
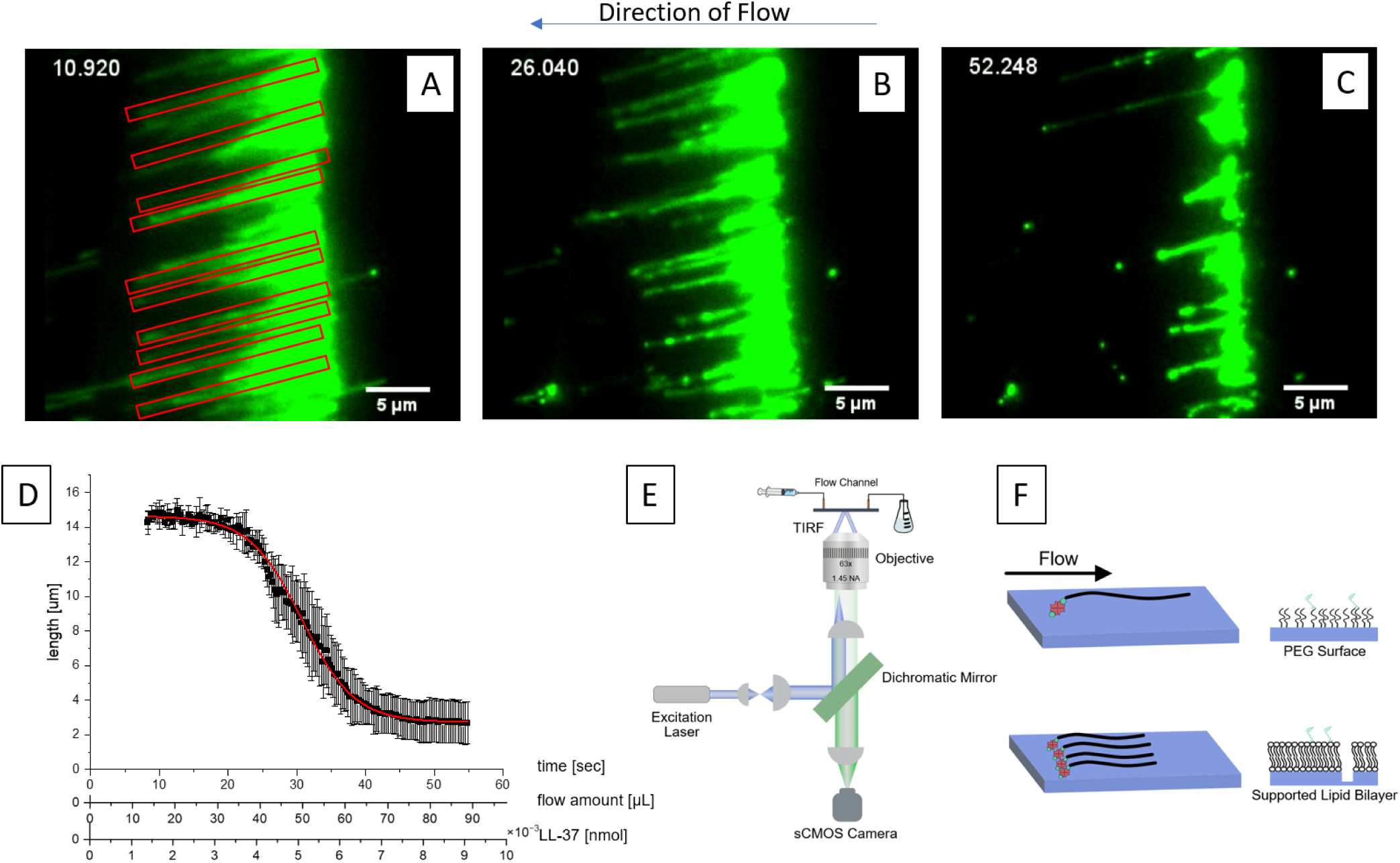
A: Biotinylated λDNA was selectively attached to a glass slide with lipid bilayers to form so-called DNA curtains in a flow channel. λDNA was fluorescently labeled and imaged using high resolution fluorescence microscopy. λDNA strands in curtains are stretched out due to shear forces of the applied flow and condense into dense structures under the influence of 100 nM LL-37 (B and C). A higher LL-37 concentration was chosen for this experiment to compensate for the presence of an increased amount of DNA. D: Marked single molecule strands in A were used to determine the condensation constant by plotting DNA length vs time. Time can also be converted into amount of flow in μL or concentration of LL-37 in this specific system. Ten strands were analyzed, and the condensation constant was determined using a Boltzmann Sigmoid Fit. Standard deviation is shown. E: Set up of the experiment. F: Schematic of a DNA curtain on different surfaces.

However, we more often observed λ DNA condensation by LL-37, followed by immobilization onto the modified surface than fully condensed λ DNA. In **Figure S4**, we show networks of DNA forming upon binding by LL-37 within λ DNA curtains. We thought that immobilization may be due to LL-37 binding to the supported lipid bilayer. To test this hypothesis, we used PEG modified surfaces to image single λ DNA molecules.^48^ **Figure 2** shows that on PEG modified surfaces, which typically serve as an inert barrier to non-specific adsorption, LL-37 condenses and immobilizes λ DNA on pre-formed complexes. This result indicates that upon binding, LL-37 changes the surface properties of the negatively charged polymer, causing condensation with itself and the nearby surface.

Because LL-37 shows oligomerization with itself in solution^49^, we tested the effect of peptide concentration on λ DNA condensation and immobilization. We varied the concentration of LL-37 in experiments using both λ DNA curtains and PEG-modified surface to see if condensation was concentration dependent. Indeed, at or below 10 nM LL-37 we observe little or no condensation of λ DNA. However, at 25 or 100 nM we observe rapid condensation as well as immobilization, indicating a concentration dependent effect on λ DNA binding.

To test whether LL-37 is capable of fully condensing λ DNA in solution, we perfomed experiments by pre-mixing the peptide with λ DNA, then adding the complexes into the flow cell, followed by washing the channel with 1X PBS and SYTOX orange and observing the formed complexes. Below 10 nM LL37, we observe little or no DNA attachment to the flow cell surface (**Figure S5A** and **S5B**, **Supplemental Movie S1** and **S2**). At concentrations of 100 nM LL-37, we observe condensation of and partially immobilization of region along λ DNA molecules, with some regions along the DNA chain that diffuse freely (**Figure S5C, Supplemental Movie S3**). These observations indicate that in solution LL-37 binds λ DNA, and condenses the dsDNA molecule in conditions where λ DNA is relaxed, rather than stretched out by flow.

### All λ DNA in a system is complexed by LL-37 at a (w/w) ratio of 1:1.7

The previous experiments show the substantial impact of LL-37 peptide on λ DNA conformation and LL-37’s condensation capacities. However, it is difficult to draw quantitative conclusions about condensation from these experiments due to varying amounts of nucleic acid attached to the surface, and hence varying amounts present in the system as a whole. Additionally, active flows in the system where λ DNA is stretched out influence the interaction of proteins with λ DNA,^50^ also considering that the nucleic acid is attached at one end to the surface. Hence, subsequent experiments were performed with complexes prepared in vials.

Electrophoretic Mobility Shift Assays (EMSA) using agarose gels were performed to investigate complex formation of different (w/w) ratios of both biomolecules (**Figure 3**). At a w/w ratio of λ DNA/LL-37 of 1:1.5, a second band appears at shorter migration times, indicating the formation of a complex with larger size and molar mass, with a band showing remaining free λ DNA present in the system. At a ratio of λ DNA/LL-37 of 1:1.7, there was no band indicating free λ DNA, implying that all λ DNA was in complexation with LL-37 and bound and condensed. A formed complex having shorter migration time agrees with ζ-potential measurements of the complexes (**Figure S1**). At higher LL-37 ratios, the complexes show a reduced ζ-potential, which further explains the shorter migration distance on the shift assays.

**Figure 3.**
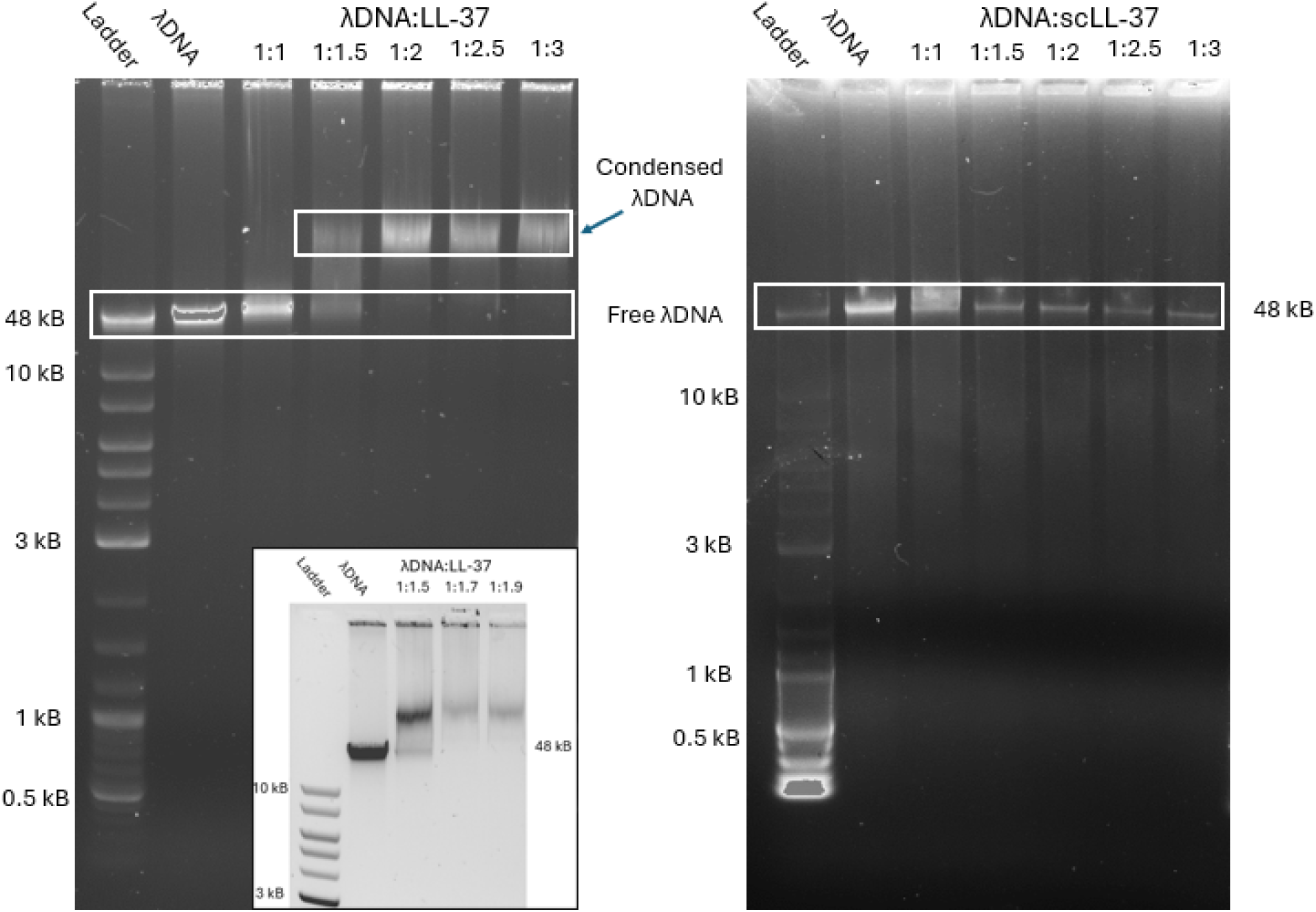
Left: Electrophoretic Mobility Shift Assays using agarose gels for investigating binding of λDNA to LL-37 at different (w/w) ratios. At a ratio of λDNA/LL-37 of 1:1.5, a larger complex is formed, indicated by the shorter migration time through the gel. At higher ratios, all DNA is in complexation and no free DNA is present any longer. Inset: Ratios between λDNA/LL-37 of 1:1.5 to 1:1.9 show that all free DNA is bound at a ratio of 1:1.7. Right: Assay using the same conditions and ratios but with scLL-37 shows that no complexes are formed.

### LL-37’s α-helical structure is essential for nucleic acid binding

Interestingly, control experiments for all three abovementioned experiments using the scrambled-sequence LL-37 negative control (scLL-37, see Supplemental Material for differences between LL-37 and scLL-37) instead of LL-37, did <u>not</u> show any complex formation. EMSA showed bands consistent with the native λ DNA band, but no bands were found at higher molar mass for the ratios that showed λ DNA/LL-37 complexation (**Figure 3**). For the microfluidic flow cells with either single molecules or DNA curtains, no condensation patterns or other interactions were observed (**Supplemental Movie S4**). scLL-37 has the same molar mass, net charge, and amino acids as LL-37, but has a different arrangement of the amino acids, which results in the loss of the α-helical structure of LL-37 and its discrete hydrophobic and cationic regions. The experiment indicates that the presumably dominant electrostatic interactions between the positively charged LL-37 and the negatively charged backbone of the nucleic acid are not the main driving force for the complexation; an amphipathic helical structure is required. Since we determined that scLL-37 does not interact with λ DNA, subsequent experiments were performed with LL-37 only.

### LL-37 blocks and protects the DNA’s phosphodiester backbone and binds in the minor groove of A-T rich sequences of λ DNA

Agarose gel assays of complexes treated with DNase I and the protease trypsin (**Figure S2**) at high ratios of LL-37 illustrated that the complexes are resistant to degradation, which agrees with literature.^18^ Formed complexes that were incubated with DNase I at 37 °C for 2 hours do not show any signs of degradation in contrast to uncomplexed λ DNA (**Figure S2**). DNase I, a deoxyribonuclease, degrades nucleic acids via hydrolysis of their phosphodiester backbone. When complexed with LL-37, λ DNA is protected from this degradation, suggesting that LL-37 is blocking access to the nucleic acid backbone.

Proteolytic treatment of the complexes with trypsin also did not show any degradation. Trypsin is known to cleave peptide bonds between either the carboxyl group of arginine or the carboxyl group of lysine and the amino group of the neighboring amino acid. LL-37 alone is readily degraded by this protease.^51,52^ Here, trypsin is unable to cleave these amino acids of LL-37 in the complexes, indicating that both arginine and lysine may be directly interacting with the nucleic acid. Whereas arginine features a guanidino side chain, lysine has a butylamine side chain. Both are protonated or partially protonated when at physiological pH and hence, are cationic under physiological conditions. It is very likely that these two cationic amino acids therefore interact with the anionic phosphodiester backbone of the nucleic acid.

4ʹ,6-Diamidin-2-phenylindol (DAPI) is a DNA-specific probe which forms a fluorescent complex by predominantly attaching in the minor grove of A-T rich sequences of DNA (in rare cases DAPI can intercalate with DNA molecules what is not relevant to the present study).^53^ Using a DAPI displacement assay (**Figure 4A**) emission quenching from the DAPI-λ DNA signal was observed returning back to baseline DAPI fluorescence intensity when LL-37 was added to the mixture. This decrease in fluorescence is most likely a result of displacement of the DAPI molecules by the peptide from the λ DNA A-T rich grooves. These results align with previous studies on DAPI nucleic acid complexes.^53^ After normalizing all fluorescence intensities to 1 [a.u.] a shift in the signal maximum to higher wavelengths is visible for the complex formed at a (w/w) ratio of λ DNA/LL-37 of 1:1.7 (**Figure 4B**), the ratio where no native λ DNA remains, according to previously discussed results (**Figure 3**). Without the overwhelming signal contribution of native λ DNA, the shift is visible, what may be related to the DNA condensation with LL-37.

**Figure 4.**
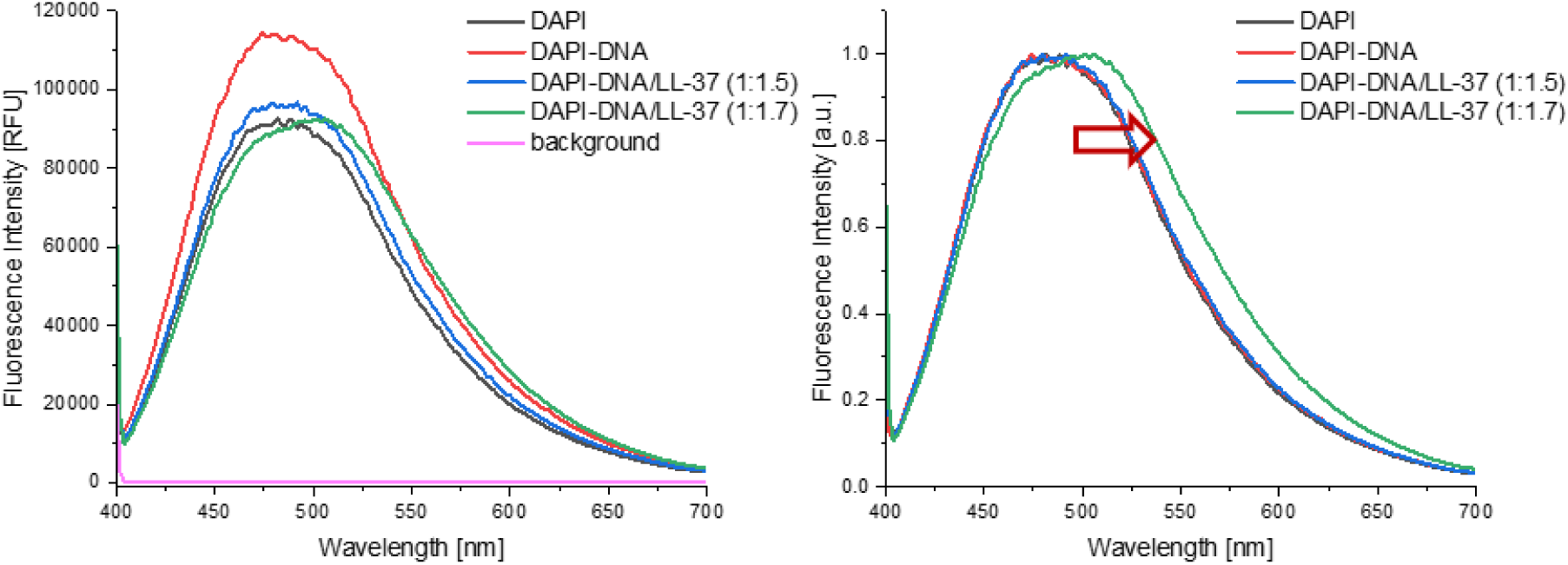
DAPI fluorescence assay to investigate binding mechanism of LL-37 and λDNA. Left: Fluorescence intensity in RFU plotted vs emission wavelength in nm. λDNA increases DAPI fluorescence intensity through interaction with the fluorescent probe. LL-37 in complexation with λDNA decreases fluorescence intensity, indicating a replacement of DAPI by LL-37. Right: Normalized fluorescence intensity in a.u. plotted vs emission wavelength in nm. λDNA in complexation with LL-37 at a (w/w) ratio of 1:1.7 shows a distinct shift of the maximum to higher wavelength, indicating further the replacement of DAPI when λDNA is in complexation with LL-37.

### LL-37 complexation of λ DNA results in loss of distinct DNA conformation

Circular dichroism (CD) measurements of the complexes to observe changes in conformation of λ DNA showed that indeed the nucleic acid loses its random coil structure. CD analysis of λ DNA only and complexes at different (w/w) ratios are shown in **Figure 5**. Λ DNA alone shows distinct B-form features with maxima at 220 and 270 nm and a minimum at about 245 nm, matching reports in the literature for nucleic acids.^54,55^ Upon the addition of LL-37, the maxima and the minimum subduct to lower intensities. At a (w/w) ratio of 1:1, the maximum at 220 nm disappeared, while at the 1:2 ratio, all three distinct features of the CD spectra disappeared. LL-37’s CD spectra shows it is α-helical, as expected, with two minima at 208 and 225 nm, a maximum at about 215 nm, and with no signals above 250 nm.^4^ With increasing peptide content in the complex, the analyzed complex starts to resemble more and more a sample of LL-37 only in conformation. By comparing the complexation ratio of λ DNA/LL-37 of 1:2.5 (**Figure 5**, cyan) to LL-37 only (magenta), which is analyzed here at the same concentration but without λ DNA, it is apparent that the spectrum represents a signal of LL-37 only. With excess of the peptide, the complex seems to precipitate out and not be detectable by CD, leaving only excess LL-37 to be analyzed in solution.

**Figure 5.**
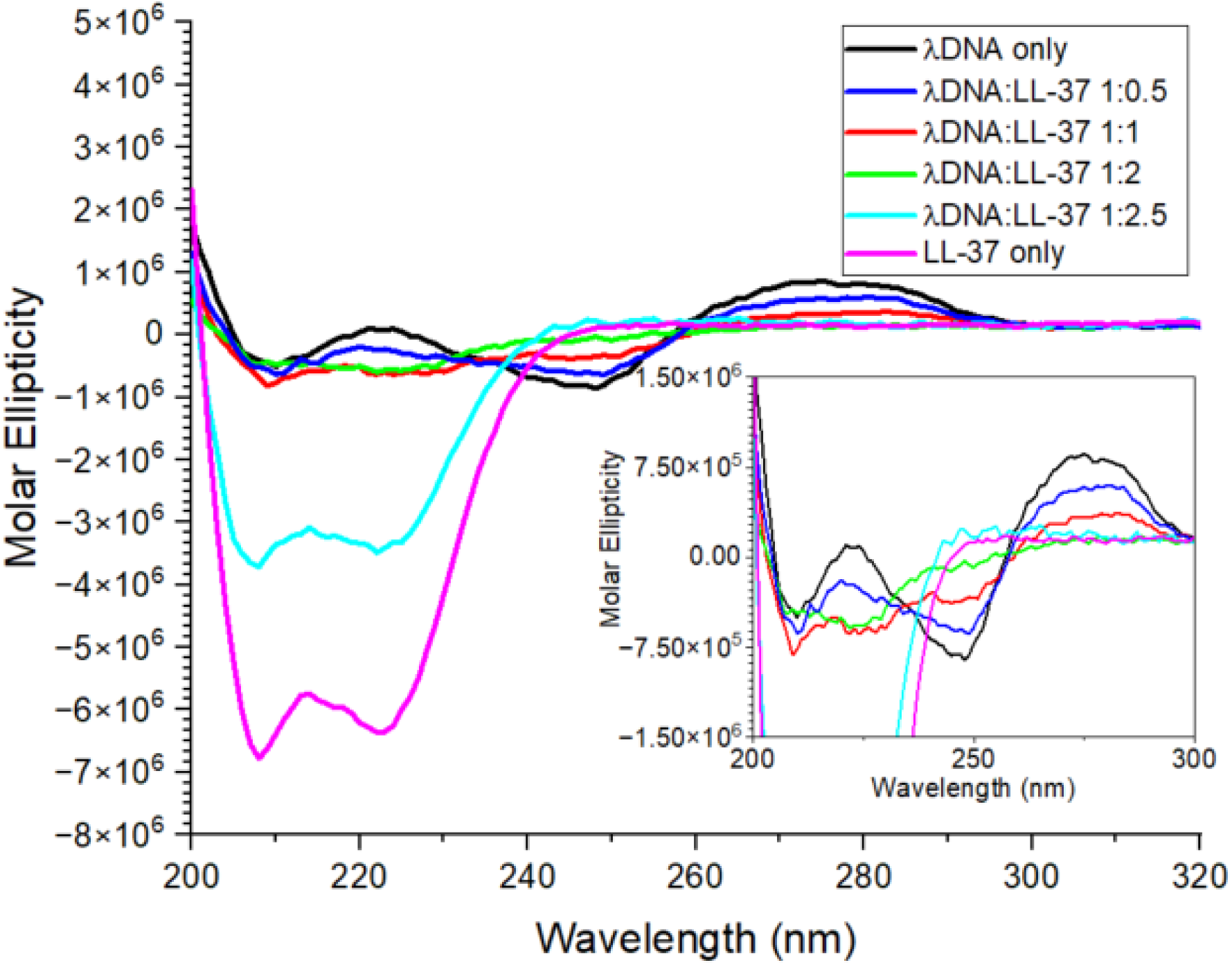
Circular Dichroism measurements of λDNA to LL-37 at different (w/w) ratios. The loss of the distinct DNA signal is visible as more LL-37 is added.

### Atomic force microscopy illustrates concentration dependency of complex conformation

To image the complexes at different ratios and to investigate their size, atomic force microscopy (AFM) in an aqueous PBS buffer matrix was employed. AFM is a high-resolution non-optical imaging technique that uses optical feedback from a sharp AFM tip connected to a moving cantilever that scans over a sample surface to obtain lateral and vertical sample information.^56,57^ Complexes were formed in vials at different ratios prior to transfer onto the mica surfaces for analysis. **Figure 6** shows the micrographs and their respective height profiles. At a ratio of 1:2.5 of λ DNA to LL-37—simulating an excess of LL-37 in a system—complexes show circular structures with an approximate diameter of 150 nm and a height of about 10-15 nm, indicating a disc-like condensate shape. With decreasing amounts of LL-37, the complexes appear to become smaller while maintaining the disc-like structure (1:2 (w/w) λ DNA/LL-37), before becoming less uniform (1:1 (w/w) λ DNA/LL-37) and, finally, with an excess amount of λ DNA present, small nodes are formed with many DNA strands loosely arranged around the node.

**Figure 6.**
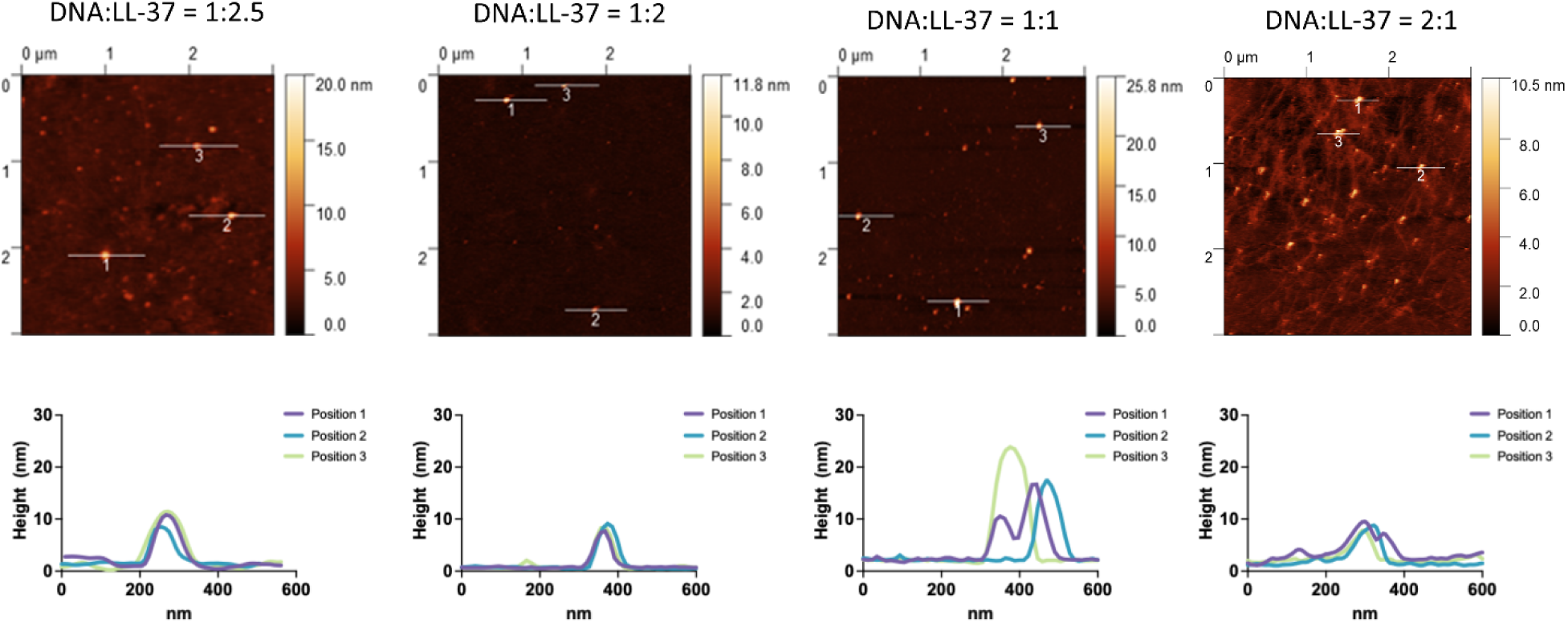
Atomic Force Microscopy of LL-37 in complexation with λDNA at different w/w ratios. At estimated full complexation, complexes show a diameter of about 100-150 nm and a height of 10 – 15 nm. With decreasing LL-37 concentration (left to right), complexes appear larger and less defined, with free λDNA clearly evident in the rightmost panel.

### AF4-MALS and SAXS confirm disc-like complex structures at LL-37 overexpression levels

To confirm the size of the complex at excess concentrations of the peptide, we employed asymmetrical flow field-flow fractionation in combination with a multi-angle light scattering and a UV/Vis detector (AF4-MALS-UV), a technique previously shown to be appropriate for molecular interaction studies of peptides.^58–60^ Due to the experimental nature of the technique, it was not possible to achieve separation of the complexes in normal mode, as the focusing step caused the analyte to elute in the void peak **(Figure S3**). Generally speaking, normal mode AF4 should be able to separate particles of 150 nm in diameter, and elution in the void peak without separation is a sign of very large-sized analytes. The elution mode of the AF4 separation was therefore changed to steric mode. ^61^ Results showed indeed r_rms_ of 75 nm (**Figure 7A**, for ease of comparison plotted as diameter of gyration Dg) with a polydispersity index of 2.15, which confirms the AFM results with a diameter of 150 nm. However, due to the steric nature of the elution, no further evaluation on the conformation, such as r_rms_/r_h_ shape factors or Kratky plots was possible^62–64^. Therefore, and to confirm the disc-like structure, small angle X-ray scattering (SAXS) data were obtained as shown in **Figure 7B**. A distinct differentiation between the calculated average of LL-37 and λ DNA alone and the data of the complex is visible, whereas a simulation of a disc-like structure of 150 nm in diameter and 15 nm in height with added polydispersity is in agreement with the data of the λ DNA/LL-37 complex (see Supplemental Material).

**Figure 7.**
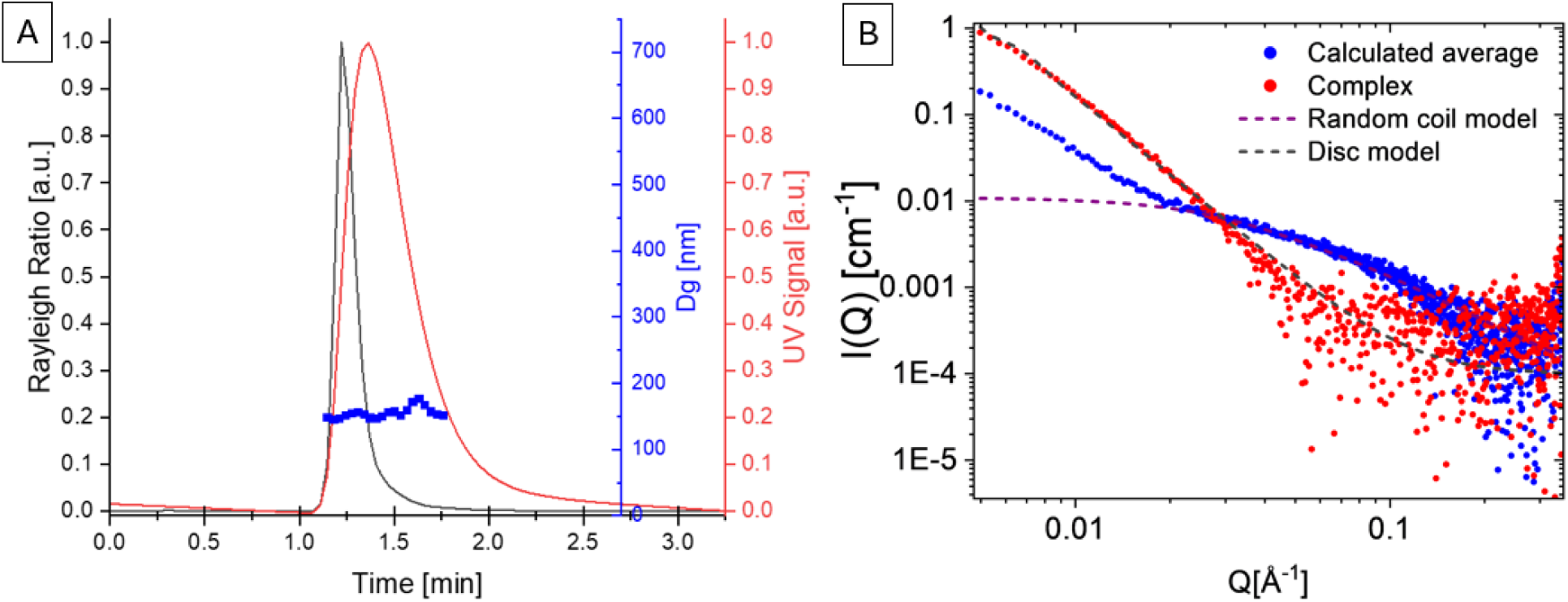
A: AF4-MALS-UV in steric elution mode of complexes at a ratio of λDNA/LL-37 of 1:2.3 displays structures with a diameter of about 150 nm and a polydispersity index of 2.15. Rayleigh ratio of the MALS detector (black, a.u.), UV signal (red, a.u.) and obtained diameter of gyration Dg (blue, nm) plotted vs retention time. B: SAXS data for the calculated combined average of λDNA and LL-37 alone (blue) and of the complex at a λDNA/LL-37 ratio of 1:2.3 (red) plotted alongside the simulated data for a disc-like structure with a 150 nm diameter, 15 nm height and added polydispersity.

### LL-37 compacts DNA in Neutrophil Extracellular Traps in a dose-dependent manner

Having shown that LL-37 has an impact on nucleic acid conformation and mobility, we next investigated the impact of the peptide on neutrophil extracellular traps (NETs). The major part of the NET composition is nucleic acids, and LL-37 is naturally produced by neutrophils; however, upregulation of LL-37 due to inflammatory signals caused by the extended presence of uncleared or poorly condensed NETs may have an effect on their structures over time.

For initial evaluation, fluorescently labeled *in vitro* NETs for nucleic acid visualization were treated with 100X the physiological concentration of LL-37 (115 μg/mL) found in human blood plasma (1.15 μg/mL)^65^ and imaged using high resolution confocal microscopy. **Figure 8A** shows time-resolved micrographs of several NETs, where multiple cells expelled their DNA to form a wide-spread network. Extracellular LL-37 is added to the cell culture (t=0 s), with an endpoint at t=5.87 min. Already after t=0.45 min, we observed an increase in the fluorescence intensity of the nucleic acid stain suggesting local DNA condensation. Compared to t=0 s, NETs appear to occupy less area at the endpoint (t=5.87 min), and exhibit a further increase in local fluorescence intensity, indicating that nucleic acid strands are compacting together.

**Figure 8.**
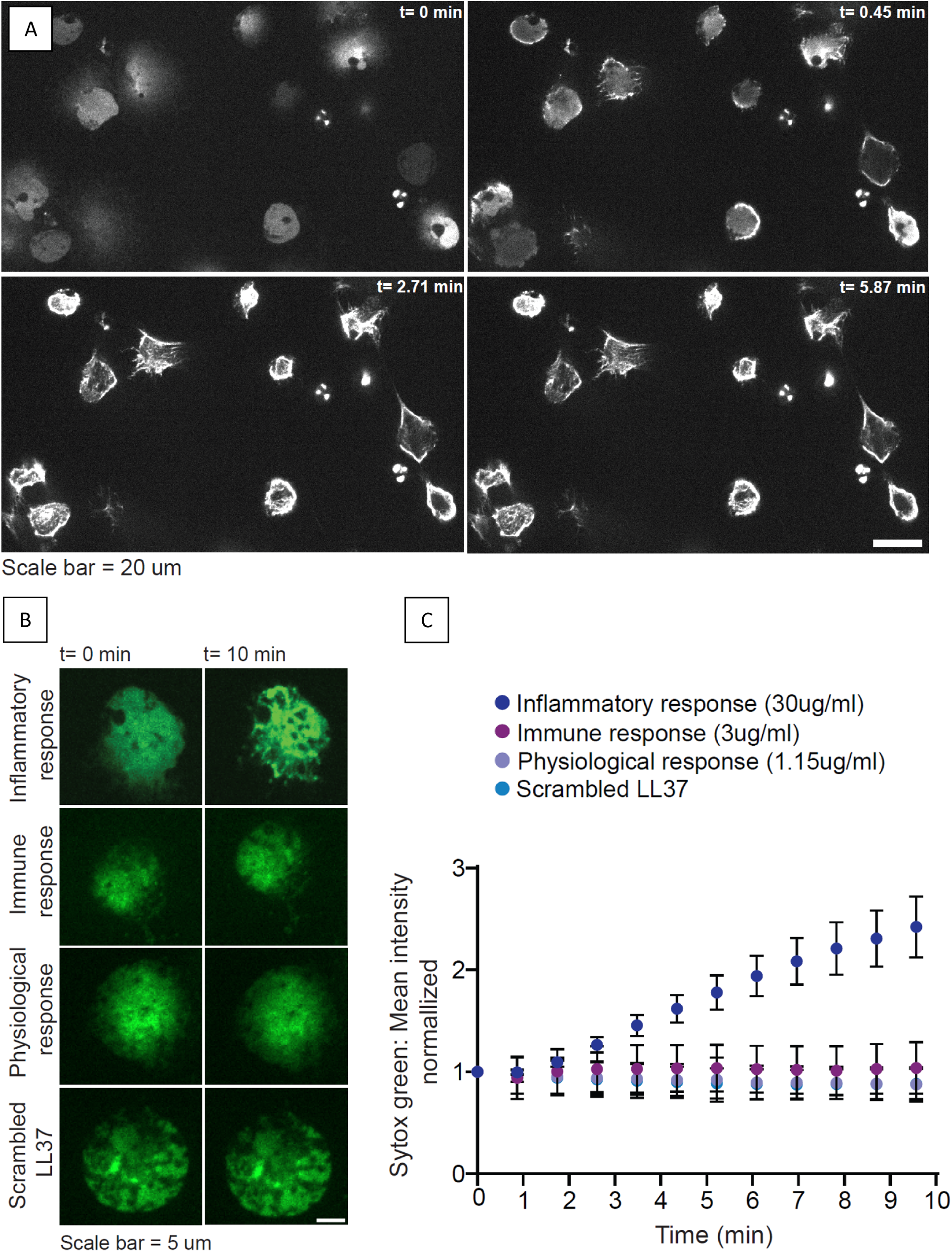
A: Example of Neutrophil Extracellular Traps (NETs) and the influence of LL-37 at 100X physiological concentration. Nucleic acids were fluorescently labeled with SYTOX Green. Compaction of nucleic acids occurs immediately after addition of LL-37, visible through the reduction in NET area. B: Representative images of NETs and subsequent compaction of nucleic acid under the influence of LL-37 at different concentrations on a single cell level, displayed at t=0s vs t=10 mins. Green fluorescence represents SYTOX Green staining of the NETs. The duration of the video was 10 min in total. Images are at coverslip-cell interface Z = 0 µm. C: Quantification of SYTOX Green mean fluorescence intensity plotted vs time after treatment with different concentrations of LL-37. Total fluorescence integrated density of SYTOX Green was normalized to the change in area over time to obtain total mean SYTOX Green fluorescence intensity. The total mean fluorescence intensity was further normalized to t=0 to compare changes in fluorescence intensity over time relative to the initial measurement. Each data point represents n=50 cells.

Here, we observed the formation of a broader interaction pattern involving several nucleic acid strands, suggesting that nucleic acids are not condensing into distinct complexes, most likely due to restrictions on nucleic acid mobility from being intertwined within the existing network, comparable to the above-described experiment using “DNA curtains”.

Subsequently, the concentration-dependency of LL-37 on NET compaction was investigated by adding extracellular LL-37 to NETs at presumably physiologically relevant concentrations. As no values of these LL-37 concentrations during NET formation in blood and/or serum have been reported in the literature, comparable findings to estimate potential concentrations for immune responses and pathological values of LL-37 were used: amounts found in healthy patients (∼1.15 μg/mL)^65^; amounts representing an immune response state (3 μg/mL)^66^; and inflammatory/pathological values of LL-37 (30 μg/mL)^67–69^, with the latter simulating severe local overexpression of LL-37. Using these conditions, we recorded time-lapse confocal movies of LL-37 treated NETs and measured the temporal evolution of the fluorescence DNA signal in the NET area.

Representative micrographs of initial and end points of LL-37 treated NETs from a single neutrophil are shown in **Figure 8B**. For these experiments, end points were reached after 130 s (figure not shown), with videos having been recorded for 10 min. We found that added inflammatory/pathological values of LL-37 significantly increase the fluorescence intensity of the nuclei acid stain on NETs, suggesting NETs compaction (**Figure 8B, C**). Physiological concentrations of LL-37, concentrations representative of an immune response, as well as scLL-37, did not show any impact on occupied NETs area and did not compact the extracellular nucleic acids, confirming earlier discussed results on DNA condensation. The results show that excess amounts of LL-37, at concentrations reported at sites of acute inflammation, compact NETs acids in a dose- and time-dependent manner, suggesting an impact on NETs fate or function during acute inflammation.

## 3. Discussion

With the study presented herein, we attempted to further investigate the interactions of the human cathelicidin LL-37 with nucleic acids, as the formed complexes have been shown to have both beneficial and detrimental effects on human health.^17^ Using λ DNA, we gained more insights into the binding mechanisms of LL-37 and believe that there are interesting implications for the roles that LL-37/DNA complexes may play in the human immune response in health and disease.

The scrambled-sequence version of LL-37, while consisting of the same amino acids as LL-37, does not form an α-helical structure, and was previously shown to not possess the same antibacterial properties as LL-37.^70^ Interestingly, a reversed LL-37 molecule with the intact α-helical structure showed comparable functions to native LL-37, indicating the importance of the structure and amphipathic nature of the peptide.^71^ In several of our experiments (See **Figures 3 and 8**), no interactions between scLL-37 and λ DNA were observed at the investigated ratios. Given these results, electrostatic interaction as the main contributor for complexation can likely be excluded. The α-helical structure of LL-37 results in one face being very hydrophobic, while the other is positively charged. Therefore, LL-37 is most likely binding non-specifically to dsDNA via the negatively charged phosphate backbone. This binding then results in dimer or tetramer formation, causing DNA to condense or immobilize on the surface if the hydrophobic face is exposed to solution. scLL-37 has the same charges, but they are not presented on secondary structure to allow for non-specific binding to dsDNA the same way as LL-37. The strong interaction of dsDNA strands after electrostatic contact with LL-37 is therefore believed to be due to hydrophobic or aromatic interactions with LL-37 molecules, which contain four phenylalanines. Attraction between the cationic region of LL-37 to the negatively charged backbone of the nucleic acid molecule likely exposes the hydrophobic region, causing interactions with other hydrophobic moieties close by, such as the DNA’s nitrogenous bases.^72^ These interactions would further distort the classic B form of the nucleic acid.^73^ An alternative theory could be that the positively charged side of LL-37 (with arginine and lysine) does bind electrostatically to the negatively charged backbone of dsDNA. The hydrophobic face of the LL-37 α-helix is then exposed to either bind to the surface as in our flow cell experiments or form multi-mers, bringing DNA strands together to condense DNA. Most likely, all of these different interactions play a dynamic role in complex formation, stability, and complex condensation over time, given that sufficient relative amounts of LL-37 are present.

As also seen in our CD results (**Figure 5**), it was previously shown that dsDNA, such as λ DNA in its classic double-stranded B form helix, does indeed interact with peptides causing a loss of normal conformation for the nucleic acid, and indicating that the nitrogenous bases are involved in the binding. Interaction and binding can be present both in the nucleic acid’s major and minor grooves.^74,75^ Binding occurs either through the peptide recognizing a specific pattern of DNA bases—the so-called base readout—or through recognizing sequence-dependent nucleic acid shapes—the shape readout.^76^ Here, shape describes structural features of the nucleic acid binding sites, with the binding mechanism being a match between groove width and electrostatic potential. A shape readout in the minor groove is usually possible through the recognition of the geometry through positively charged amino acids in the peptide, such as the arginine and lysine side chains in LL-37 (see Results Section above) forming cations at physiological pH.^77^ However, not all peptides binding to DNA will show condensation and subsequent loss of the B form.^77^

Hydrophobic interactions may also be involved in the DNA-to-lipid bilayer interaction when LL-37 is present. Lipid bilayers were used to create DNA curtains (**Figure 2**), and in situations where the barrier was neither deep nor wide enough for the nucleic acid strands to move freely above the barrier, the nucleic acid showed rigidification when under LL-37’s influence instead of condensation (see **Figure S4**). Interactions of LL-37 with lipid bilayers^16^ and other biomolecules^12,14,17^ have been shown before, and, especially when considering physiological processes or human innate immunity, these phenomena should be kept in mind. In other words, the interaction between LL-37 and nucleic acids is not occurring in an isolated system, but rather in the presence and under the influence of other biomolecules.

The upregulation of LL-37 has been previously suggested to improve human health,^78–80^ and this LL-37 regulation may be influenced by the conformation size and shape of complexes. We observed the presence of loosely arranged nucleic acids around a node with low levels of LL-37 with more free DNA visible, indicating that the full condensation of nucleic acids by LL-37 is indeed quite concentration-dependent (**Figure 6**). Underexpression of LL-37 may yield LL-37/DNA complex structures comparable to results found by Lande *et al.*, for complexes extracted from psoriatic skin lesions.^32^ We hypothesize that this could indicate that an underexpression of LL-37 (as opposed to overexpression, as implied in the past) may be involved in human autoimmune diseases such as plaque psoriasis. As we show, the structure of these complexes changes with increasing amounts of LL-37, finally forming compact disc-like structures at high levels of LL-37. Such well-compacted LL-37/DNA structures would more easily phagocytosed by macrophages.

Comparing with the published literature, the AFM observed disc-like structures that have not been observed before for LL-37/DNA complexes (**Figure 6**), but the ability of λ DNA to form disc-like structures (such as toroids, or donut-shaped structures) under the influence of condensing agents has been discussed previously.^81,82^ However, to exclude the possibility that the results are an artifact of the AFM surface encounter technique with a tip scanning over the sample, subsequent experiments were needed to confirm the disc-like structures in solution.

We employed AF4 in combination with multiple detectors and synchrotron SAXS to confirm the size and conformation of the disc-like structures. During normal elution mode of the fractionation technique, no separation was achieved and the analyte eluted in the void peak. The presence of charges—as confirmed for the complexes through ζ-potential analysis (**Figure S1**)—can cause more highly aggregated structures due to the increased concentration profile during the techniques’ focusing step, or interactions with the ultrafiltration membrane.^83^ To avoid these interactions, the use of surfactants is typically recommended for the AF4 analysis.^83^ As surfactants will change the binding pattern of biomolecules, this option was not pursued for the present study, and elution in steric mode was performed instead. Using a combination of AF4-MALS-UV and SAXS, it was possible to confirm the existence of the observed disc-like structures, and to exclude the possibility of the structures being an artifact of the AFM approach.

Other shapes that have previously been reported in literature are, for example, liquid-crystalline ordered columnar square lattices^31^, and globular structures ^46^. Specifically relevant is a study by Lee *et al.*, where complexes were also formed between LL-37 and monodisperse λ DNA.^30^ There, resulting LL-37/DNA complexes were suggested to consist of cationic protofibril LL-37 assemblies that interact with anionic DNA to form a two-dimensional columnar square lattice, which then presented ordered nucleic acid scaffolds to TLR9^30^. Structure analysis in that study was performed by SAXS, and concentrations of λ DNA were described as 1-5 mg/mL mixed with the peptide at different charge ratios. When mixing λ DNA at a concentration of 1 mg/mL with LL-37 at a (w/w) ratio of 1:2.5, our group observed phase separation, in apparent agreement with the authors’ observation, as thorough mixing, centrifugation and precipitation is mentioned for sample preparation, indicating that they also observed phase separation. For our reported results, these samples were not evaluated, as the high concentrations did not seem to be physiologically relevant, given that human LL-37 levels in different systems are reported to range from 1.15 µg/mL to 30 µg/mL^67–69^. Nevertheless, the study by Lee *et al.*, gives insights into potential therapeutical approaches and demonstrated that these complexes can trigger strong recognition by TLR9 due to their spatially-periodic DNA organization, and hence, a structure-dependent reaction was observed.^30^ Especially when discussing activation of certain receptors relevant for human health, the structure of the ligand is of outmost importance. In general, different TLRs will recognize different surface and/or intracellular components of microorganisms or molecules, and their interactions trigger the activation of the innate immune system, as well as the development of acquired immunity.^17,84,85^ Our results demonstrate the importance of peptide amounts in a system for the formation of differently shaped complex structures, which might then in turn have different impacts on human innate immunity.

In addition, we showed that the concentration of LL-37 is also relevant for the compaction of nucleic acids in NETs. Our results indicate that concentrations of LL-37 at levels of inflammatory response levels—usually triggered through a strong immune reaction—compact nucleic acids to effectively greatly reduce NET area. A reduction in NET area may be beneficial for their clearance: employed macrophages will need to cover a smaller area during phagocytosis and experience a reduced risk of getting entangled in the NETs themselves.^86^ Increased levels of LL-37 also differentiate monocytes into MΦ-1 macrophages, which in turn are more effective in NET clearance than MΦ-2 which result from NET presence during differentiation^87,88^. This switch in differentiation of macrophages might help prevent NET-related diseases and autoimmune reactions such as thrombosis. On the other hand, the other commonly used NET clearing mechanism through DNase I degradation may be hindered by compaction with LL-37, based on the discussion above regarding LL-37 and nucleic acid complex stability. Furthermore, literature findings showed previously that LL-37-treated NETs were distinctly more resistant to *S. aureus* nuclease degradation than nontreated NETs.^89^ If compaction through LL-37 increases resistance to degradation, accumulation of NETs might follow, what was, for example, found to provide an immunosuppressive microenvironment favoring the survival of premalignant cells and cancer cells, causing tumor growth.^90^ To thoroughly investigate the impact that LL-37 upregulation might have on extracellular traps and the resulting health effects, further studies will be needed.

The present study underlines the hypothesis that the amount of LL-37 available in a system for complex formation with nucleic acids is an important factor for the size and structure of formed complexes, and that expression levels of LL-37 in the human body may have an impact on human health. As discussed, different shapes have been previously reported for complexes formed between the antimicrobial peptide and nucleic acids. The presented results indicate that LL-37 at lower concentrations forms loosely aggregated DNA structures, while an excess of the peptide in the system creates dense structures. These differences in complex structure may be the reason for varying results found in literature about the functional properties of these complexes in the human body. The observed disc-like structures when excess of the peptide is present have not been reported in literature before; however, this might also be due to the differing sizes of nucleic acids. Here, λ DNA was used due to its known sequence, reproducibility and uniformity. λ DNA is a large phage-derived DNA molecule and might not be fully representative of nucleic acids found in the human body (*e.g.*, will have a different state of methylation). Nevertheless, these results are valuable to show the impact of a small peptide on nucleic acids. The condensation of the nucleic acids is a result of interactions of the phosphodiester backbones of the nucleic acids and the protonated side chains of arginine and lysine in the peptide. Further, LL-37 may predominantly interact with the A-T rich sequences of the nucleic acids, and even though electrostatic interactions might be assumed as the main driving force of complexation, the α-helical structure is shown to be essential for complex formation. LL-37 is also shown to reduce NET size and occupation area, which may be an important step towards deciphering the molecular biophysics that may drive NET-related autoimmune diseases. Overall, the presented results highlight the potential importance of LL-37 expression levels in the human body, and its key impact on human innate immunity, with an overall implication that higher expression levels are likely beneficial to full condensation and a proper, more efficient clearance of neutrophil NETs.

## 4. Methods

LL-37 human antimicrobial peptide and its scrambled version (scLL-37) as negative control were purchased from Anaspec (Fremont, CA). Lambda DNA (SD0021) and DNase I (EN0521) were purchased from Thermo Fisher Scientific (Waltham, MA). Trypsin from porcine pancreas (T0303) was purchased from Sigma Aldrich (St. Louis, MO). Gibco PBS at pH 7.4 was purchased from Thermo Fisher Scientific Inc., and PBS for AF4 running buffer was prepared according to the Gibco recipe using sodium chloride (Thermo Fisher), sodium phosphate dibasic heptahydrate (Sigma) and potassium phosphate monobasic (Sigma). Other method specific materials are mentioned in the methods section.

To avoid adverse interactions due to storage buffer conditions, λ DNA was buffer exchanged to PBS and upconcentrated prior to experiments or complex formation, using Microspin S-400 spin columns (Cytiva) and Amicon Ultra 0.5 mL centrifugal filter (Millipore).

### Surface-functionalized flow cells for imaging single DNA molecules

Coverslips were cleaned with a 20% 7X SDS detergent solution at 90 °C for 5 min, rinsed 3 times with MilliQ water and dried with N_2_. Subsequently, coverslips were transferred into a Herrick plasma cleaner and cleaned at 100 mTorr for 3 minutes. To coat the flow cell surfaces with PEG, cover slips were reacted with APTES prior to reacting with a 2 %(w/w) solution of bio-PEG-5000-SVA and mPEG-5000-SVA (Laysan Bio, Inc) in 100 mM bicarbonate buffer.^48^ The coverslips were washed with MilliQ water and stored in −20 degrees before use. Flow cells were constructed by adhering 3M VHB double sided tape (3M, 2.5 mil, 1×2 inch strips) to pre-drilled glass slides, cutting a single channel spanning the two ports, and then attaching a cleaned coverslip. Finally, cut PEEK tubing (PEEK Tubing – 1/16ʺ OD x .020ʺ ID, IDEX) was attached to the drilled holes using 5-min epoxy (Loctite). After forming channels by affixing and Stick Slide to the PEGylated cover slips, we filled each channel with 0.5 mg/mL Streptavidin (Pierce™ through ThermoFisher) in 1X PBS using a pipette attached to silicon tubing (0.8 mm ID, Bio-Rad Laboratories) and allowed it incubate for 10 minutes to allow binding the PEG-biotin. The flow cell was rinsed with buffer before 0.1 mg/mL streptavidin (Thermo Fisher #2112) was added and incubated for another 10 minutes. In the meantime, λ DNA is biotinylated by ligation to a 3ʹ-biotinylated 12-mer oligonucleotide (5ʹ-GGGCGGCGACCT-3ʹ or 5ʹ-AGGTCGCCGCCC-3ʹ) that is complementary to one of the cohesive ends of λ DNA ^91^. After a final flushing step of the flow cell with PBS, 2 pM biotinylated λ DNA in PBS supplemented with 40 nM SYTOX orange (Thermo Fisher #S11368) was added and incubated for 10 minutes.

### DNA curtains on supported lipid bilayers

Microfluidic channels were assembled as described above, using a cleaned coverslip etched with a diamond tipped scribe to create the barrier where the DNA curtain will assemble. DNA curtains were created according to Greene *et al*. ^47^ In brief, liposomes were formed by preparing a 0.5 mL solution of 10 mg/mL of DOPC (18:1 (Δ9-Cis) PC (DOPC) 1,2-dioleoyl-sn-glycero-3-phosphocholine (Avanti 850375, 25 mg in chloroform at 10 mg/mL)) with 0.5% DPPE-biotin (16:0 Biotinyl Cap PE 1,2-dipalmitoyl-sn-glycero-3-phosphoethanolamine-N-(cap biotinyl) (sodium salt) (Avanti 870277, powder, add chloroform to get 10 mg/mL)) in 10 mM Tris pH 7.2, 100 mM NaCl. Flow cell is washed with buffer, liposomes are injected into the device and incubated for 30 min. Subsequent washing steps, addition of streptavidin as described above, before addition of 10 pM biotinylated λ DNA in PBS supplemented with 40 nM SYTOX orange.

### TIRF Imaging of single DNA molecules

Flow cells were mounted on the microscope stage and connected to a MX Series II 2-position, 6 position PEEK switching valve (IDEX). A syringe with 1X PBS and 40 nM Sytox Orange was mounted on a multi-syringe infusion pump (KD Scientific) and connected to a manual 3-way switching valve (IDEX). The valve was then connected to the switching valve using PEEK tubing. Finally, a solution containing varying concentrations of LL-37 in 1X PBS with 40 nM Sytox Orange was inject into a 0.5 mL sample loop (Cytiva) connected to the MX switching valve. For experiments, imaging acquisition was performed on Zen software. Flow rates were set to 100 µL/min on the infusion pump to extend DNA during imaging. After imaging extended DNA, the MX switching valve was switched to a position allow for injection of the LL-37 solution into the flow channel.

Single molecule fluorescence imaging was performed on an inverted Zeiss Elyra 7 microscope using a Plan-Apo 63x/1.46 NA Oil immersion objective. For Sytox Orange stained DNA, a 0.5 W Sapphire 561 nm laser was reflected into the flow channel using an MBS 405/488/561/641 filter set and focused onto the back focal plane of the objective to obtain total internal reflection. Emitted light was collected through the objective and filter set, passed through a LP 560 filter cube via a Duolink filter and imaged on pco.edge 4.2 high speed sCMOS camera. Movies were captured at an exposure time of 50 ms per frame unless otherwise noted. Images were collected at the indicated exposure times. Images and stacks were then processed and false-colored using Zeiss Zen Black, ImageJ or FIJI software.

### Electrophoretic gel mobility assays using agarose gels

Agarose LE (Roche Switzerland, #11685660001) 0.5 % gels were prepared in TAE buffer (Sigma) and stained with GelRed Nucleic Acid Stain (Millipore, SCT123). Complexes were formed with a constant λ DNA concentration of 50 μg/mL in PBS and varying (w/w) concentrations of LL-37 and given time for 30 minutes. Complexes were stained with Gel Loading Dye Purple (New England Biolabs, Ipswich, MA) and loaded at 20 μL onto the gel. A Bio-Rad PowerPac Basic (Bio-Rad Laboratories, Inc., Hercules, CA) was run for 70 min at 90 V before analyzing the bands. A 1 kb Plus DNA Ladder (New England Biolabs) was substituted with λ DNA to display the migration time of native λ DNA. For degradation assays, DNase I at 50 and 100 U/mL and trypsin at molar ratios of 1:40 and 1:60 were added according to the manufacturer recommendation to complexes before incubation for 2 hours at 37 °C and loading onto the agarose gel.

### DAPI fluorescence assay

4ʹ,6-Diamidine-2ʹ-phenylindole dihydrochloride (DAPI) was obtained from Roche Switzerland (#10236276001). λ DNA at concentrations of 50 μg/mL in PBS supplemented with 1 μmol DAPI was mixed with LL-37 at different (w/w) ratios. Samples were analyzed in 96 well plates using a Tecan Spark plate reader (Tecan Trading AG, Switzerland).

### Circular dichroism

λ DNA at concentrations of 100 μg/mL in PBS was mixed with LL-37 in PBS at different (w/w) ratios. Spectra were recorded with a Jasco Circular Dichroism Spectrophotometer J-1000 (Jasco, Tokyo, Japan) between 320 and 170 nm with a resolution of 1 point per nm. Obtained data was then recalculated to molar ellipticity according to ^92,93^.

### Atomic force microscopy

An Asylum AFM MFP-3D (Asylum Research, Oxford Instruments. Santa Barbara, CA) in tapping mode under aqueous sample conditions was employed to gain insights into dimensions of the complexes. Cleaved highest grade V1 mica discs (12 mm, 50-12, Ted Pella Inc., Redding, CA) were placed on the AFM, coated with 0.001% poly-L-lysine for 5 min (PLL, Ted Pella, Pelco #18021) and washed 4 times with MilliQ water. PLL was used to change the surface charge of the mica to ensure detectability of the complexes. A 10X lower concentration than recommended was used, as PLL was previously shown to condense nucleic acids itself. ^94^ Control experiments with λ DNA alone were recorded to ensure that no condensation was initiated by PLL. Complexes were formed with λ DNA at concentrations of 50 μg/mL in PBS was mixed with LL-37 at different (w/w) ratios in vials and incubated on the mica disc for 20 min, washed 4 times with PBS and left under 50 μL PBS for analysis. SNL-10 silicon tip CN nitride levers (Bruker, Camarillo, CA) were used as cantilevers in tapping mode and instrument control as well as data collection were performed using IGOR Pro (WaveMetrics, Lake Oswego, OR, USA). Images and height profiles were analyzed using the open source data analysis software Gwyddion ^95^.

### ζ-potential analysis

Using a NanoBrook Omni (Brookhaven Instruments Corporation, Holtsville, NY) and appropriate cuvettes (Brookhaven, Fisher Scientific NC9968046), complexes at different ratios (see above) were analyzed for their ζ-potential. Standard deviation was calculated from triplicates.

### Asymmetrical flow field-flow fractionation

The asymmetrical flow field-flow fractionation (AF4) instrument used was a Postnova AF2000 MultiFlow FFF (Postnova Analytics, Salt Lake City, Utah), connected to an SPD-20A UV/Vis detector operating at a wavelength of 215 nm and a PN3621 multi-angle light scattering (MALS) detector operating at a wavelength of 532 nm (both Postnova). A PN5300 Autosampler (Postnova) handled the sample injection onto the AF4 channel. The separation channel used was a Postnova long channel with a tip-to-tip length of 30 cm, the spacer was 350 μm in height, and the ultra-filtration membrane forming the accumulation wall was regenerated cellulose with a cut-off of 10 kDa (Postnova). Performing steric elution mode experiments, 20 μL of sample were injected at a flow rate of 0.2 mL/min with a detector flow rate Q_d_ of 4 mL/min and a constant crossflow Q_c_ = 0.25 mL/min to stabilize the elution through the separation channel. Processing of the obtained data from UV and MALS detectors after separation was performed using NovaFFF AF2000 software (Postnova). The diameter d_rms_ was calculated using the Berry method^96,97^ performing a 1^st^ order fit with the data obtained from the chosen scattering angles 4–15 (28◦−124◦). For simplification, the refractive index increment dn/dc for the complex was set to an average value of 0.17 mL/g for nucleic acids^98^, while the second virial coefficient A_2_ was considered negligible.

### Small-angle X-Ray scattering (SAXS) data collection

SAXS experiments were performed at SLAC SSRL Beamline BL4-2^99^, CA, USA, with a detector distance of 2.5 meter and X-ray wavelength of λ=1.127 Å, covering a Q range of 0.005 Å^−1^ to 0.35 Å^−1^. 30 µL of sample was inserted into the flow through cell using an autosampler set up. Using a series of 12 consecutive 1 s exposures were collected for samples and buffer blanks. To maximize the exposed volume and reduce the radiation dose per exposed volume, solutions were oscillated in a stationary quartz capillary cell during data collection. Subsequently, radial integration, screening for radiation damage to the sample and scaling according to the transmitted beam intensity, as well as correction for buffer background for the obtained data was performed using an automated data reduction pipeline at the beam line. The data set was calibrated to an absolute intensity scale using water as primary standard. Data were analyzed using scattering models.

### Cell culture, NETosis assay and microscopy

Neutrophils were grown using HL-60 cells (ATCC CCL-240), cultured at 37 °C and 5% CO_2_ in RPMI 1640 media, supplemented with L-glutamine with 25 mM HEPES, 1% penicillin and streptomycin, and 15% heat-inactivated FBS. Cells were differentiated into neutrophil-like cells through addition of 1.3% of DMSO^100^. After 7 days, cells were resuspended in imaging media (RPMI 1640 lacking phenol red (Thermo Fisher Scientific Inc., 11835030), substituted with 25 mM HEPES and 1% antibiotics P/S) before imaging. dHL60 cells at a cell count of 2 x 10^5^ cells/ml were centrifuged at 2000 rpm for 5 mins, the cell pellet was washed with 200 µL of imaging media before being resuspended in 200 µL of fresh imaging media. The cell suspension was transferred into a 4-well glass bottom plate, and 200 µL of imaging media containing ionomycin at a final concentration of 4 µM was added. Subsequently, cells in plate were incubated overnight at 37 °C and 5% CO_2_.^101^ After overnight incubation, media was removed from the cells, and 2 µL of SYTOX-green stock (5 µM) in 200 µL of imaging media was added to the cells and incubated at 37 °C for 10 minutes. Subsequently, the media containing SYTOX green was removed and replaced by 100 µL of fresh imaging media. To avoid the error of diffusion, seeding of cells is performed in 100 µL imaging media only, with subsequent addition of 400 µL of LL-37 solution at different concentrations. Therefore, the final concentration of LL-37 in the experiment has to be extrapolated to a final volume of 500 µL; final concentrations were 1.15 µg/mL, 3 µg/mL, 30 µg/mL and 115 µg/mL. High resolution videos were recorded using a Nikon confocal spinning disc microscope with 60X magnification and fluorescence detection to image the NETs. Parameters for image acquisition were the following: time interval between frames = 10 sec; total duration of time-lapse = 10 min; channel for fluorescence intensity = excitation/emission for SYTOX green = 488/523 nm; Z-stack settings = coverslip-cell interface (0 µm) and 3 µm. Two frames per position were taken to establish the starting point (t=0) of the experiment. Subsequently, video recording was paused, 400 μL of LL-37 at differing concentration was added, and the recording was continued. Data analysis was performed through semi-automated image processing using ImageJ software with FIJI plugin ^102^. Changes in fluorescence intensity values over time were analyzed for the region of interest and normalized against background and occupied area. Experiments were performed in duplicates and are displayed with standard deviations of 50 cells analyzed per condition.

## Abbreviations

AF4: asymmetrical flow field-flow fractionation
AFM: atomic force microscopy
AMP: antimicrobial peptide
CD: circular dichroism
DAPI: 4ʹ,6-Diamidine-2ʹ-phenylindole dihydrochloride
DLS: dynamic light scattering
DNA: deoxyribonucleic acid
dn/dc: refractive index increment
d_rms_: root-mean square diameter
dsDNA: double-stranded DNA
HEPES: 4-(2-hydroxyethyl)-1-piperazineethanesulfonic acid
MALS: multi-angle light scattering
NETs: neutrophil extracellular traps
NK cells: natural killer cells
PBS: phosphate buffered saline
Qc: cross flow
Qd: detector flow
RNA: ribonucleic acid
SAXS: small angle X-ray scattering
scLL-37: scrambled version of LL-37
w/w: weight-to-weight ratio

## Acknowledgements

We acknowledge Dr. Florian Ermini’s help with the ζ-potential measurements, Christina Newcomb for the AFM support and training, and Karrie Weaver for facilities support at the Green Earth Sciences Building, Stanford University. The authors would like to thank Dr. Vincent Noël for accessibility and support using the SLAC Asymmetrical Flow Field-Flow Fractionation (AF4) platform. The authors would like to thank SLAC for SAXS beamtime, and Dr. Thomas Weiss and Dr. Tsutomu Matsu for support during the SAXS experiment.

## Author Contribution Statement

Claudia Zielke (CZ): acquisition, analysis, and interpretation of all data, drafting of manuscript Behzad Rad (BR): acquisition, analysis, and interpretation of microfluidic data Josefine E. Nielsen (JEN): acquisition, analysis, and interpretation of CD and SAXS data Jiaxin Li (JL): acquisition, analysis, and interpretation of electrophoretic mobility assay data Sopida Pimcharoen (SP): acquisition, analysis, and interpretation of AFM data Manasi Sawant (MS): acquisition, analysis, and interpretation of NET related data Jennifer S. Lin (JSL): coordination of experiments, drafting of the manuscript Hawa R. Thiam (HRT): conception of the work, drafting of the manuscript Annelise E. Barron (AEB): conception of the work, drafting of the manuscript

## Additional Information

The author(s) declare no competing interests.

## Data Availability

The datasets generated during and/or analyzed during the current study are available from the corresponding author on reasonable request.

