## Supplemental Information for "Investigation of Regulation and Binding Patterns of the Human Cathelicidin Peptide LL-37 in Complexation with Nucleic Acids, and its Impact on Neutrophil Extracellular Traps"

<u>LL-37 and its negative control scLL-37.</u>

| **Peptide** | **Sequence** | **Net charge** | **Molar Mass** |
|---|---|---|---|
| LL-37 | LLGDFFRKSKEKIGKEFKRIVQRIKDFLRNLVPRTES | +6 | 4493.6 g/mol |
| scLL-37 | GLKLRFEFSKIKGEFLKTPEVRFRDIKLKDNRISVQR | +6 | 4493.6 g/mol |

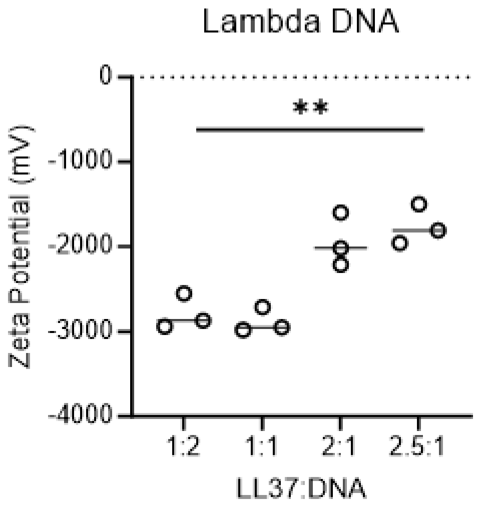


Figure S1: ζ-potential measurements of λDNA/LL-37 at different ratios. The net negative charge decreases with increasing LL-37 in complexation; however, fully condensed complexes still present a negative charge, which was confirmed through gel electrophoresis. Triplicates with average values and indicated standard deviation.

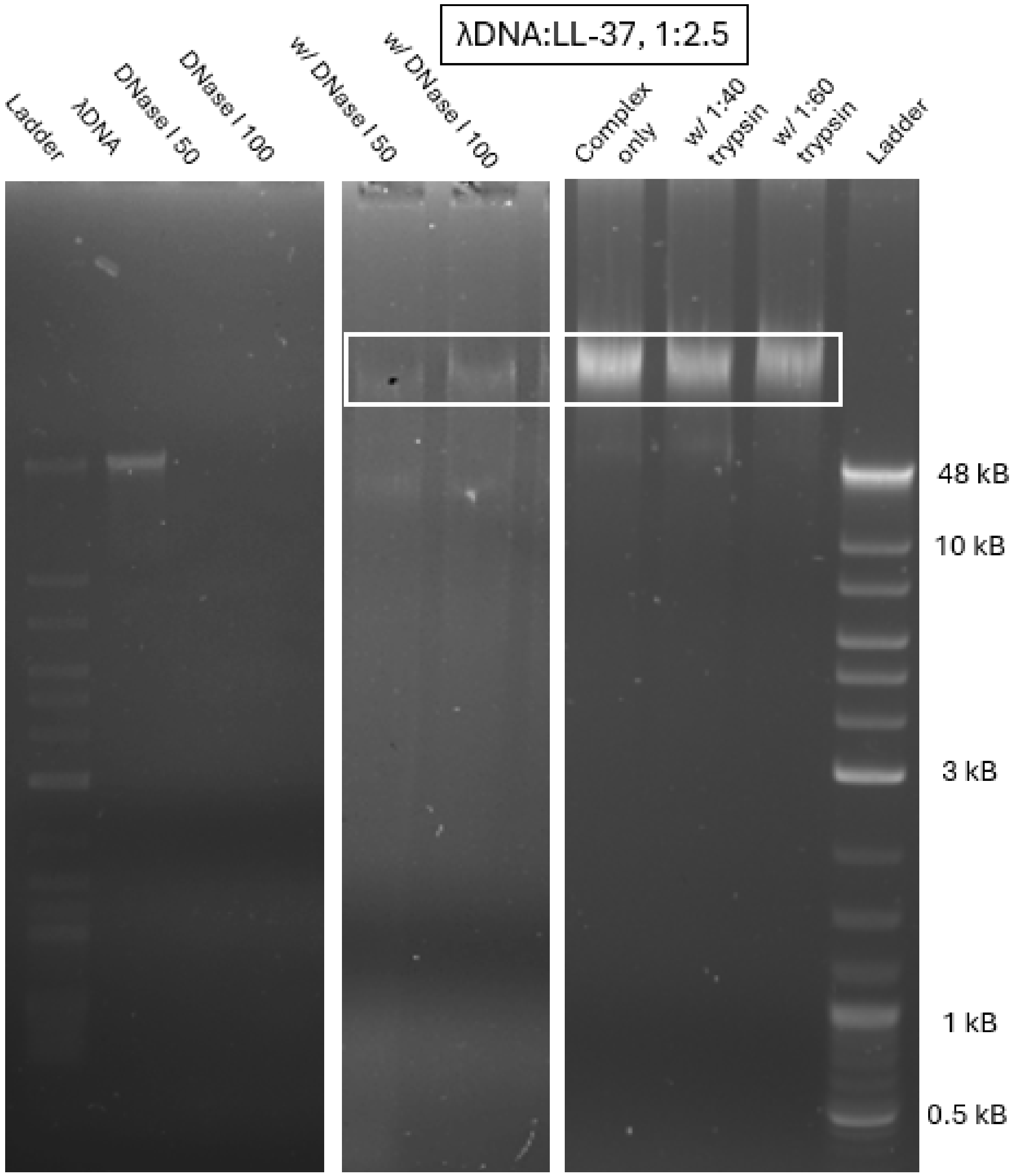


Figure S2: Electrophoretic Mobility Shift Assays using agarose gels for investigating the stability of λDNA/LL-37 complexes at a ratio of 1:2.5 following treatment with DNase I and trypsin. λDNA alone is digested by both 50 and 100 U/mL DNase I whereas in complexation, the band for the DNase I treated sample remains unchanged compared to the complex only (untreated) results. Treatment with trypsin does not digest the complexes either.

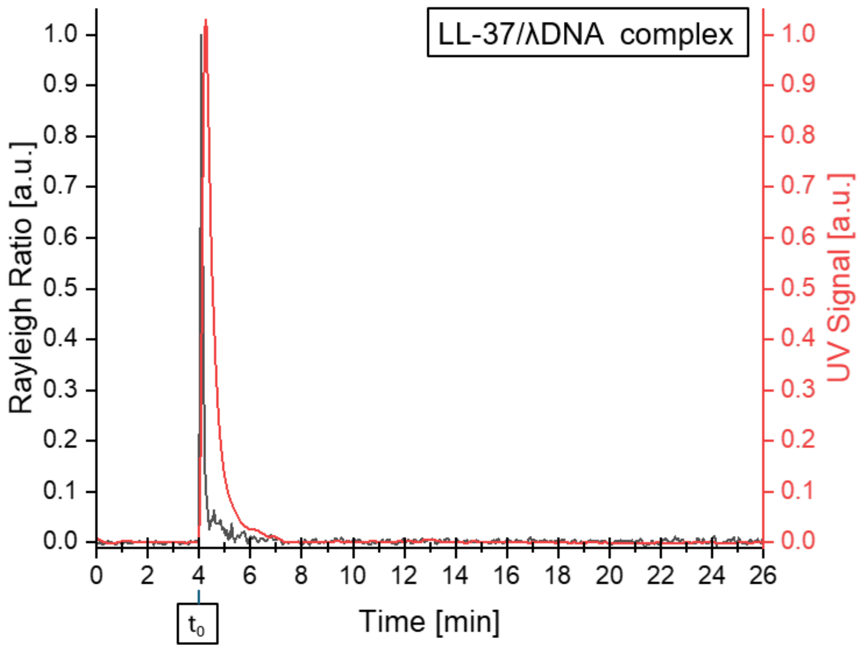


Figure S3: Example of an AF4-MALS-UV in normal elution mode of complexes at a ratio of λDNA/LL-37 of 1:2.3. Retention did not work under any varied condition, and complexes eluted in the void peak at t=0. Rayleigh ratio of the MALS detector (black, a.u.) and UV signal (red, a.u.) plotted vs retention time.

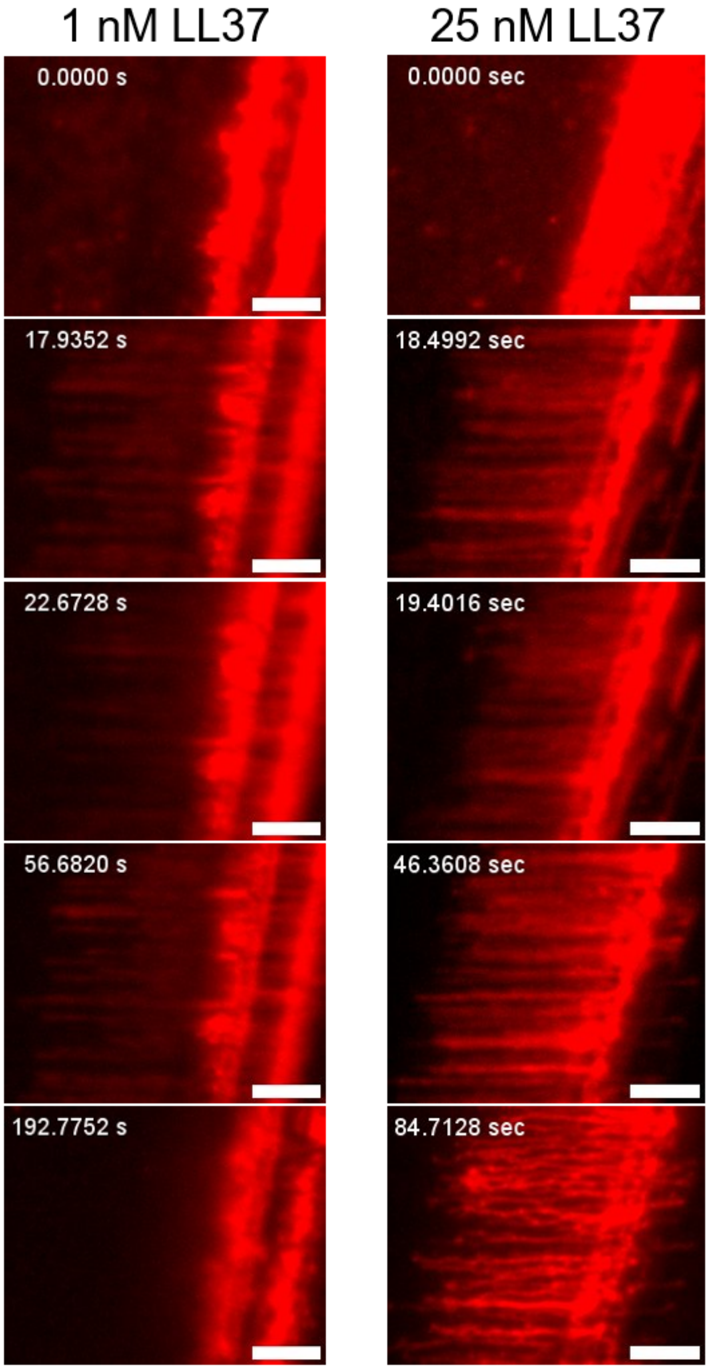


Figure S4: **LL37 condenses λDNA under flow.** Images of λDNA 'curtains' formed on a mobile supported lipid bilayer. At time 0 s, the flow is off and the DNA is not extended. Upon turning on the flow, the DNA extends towards the left. LL37 is introduced starting at ~20 seconds in each experiment. Scale bars are 5 µm.

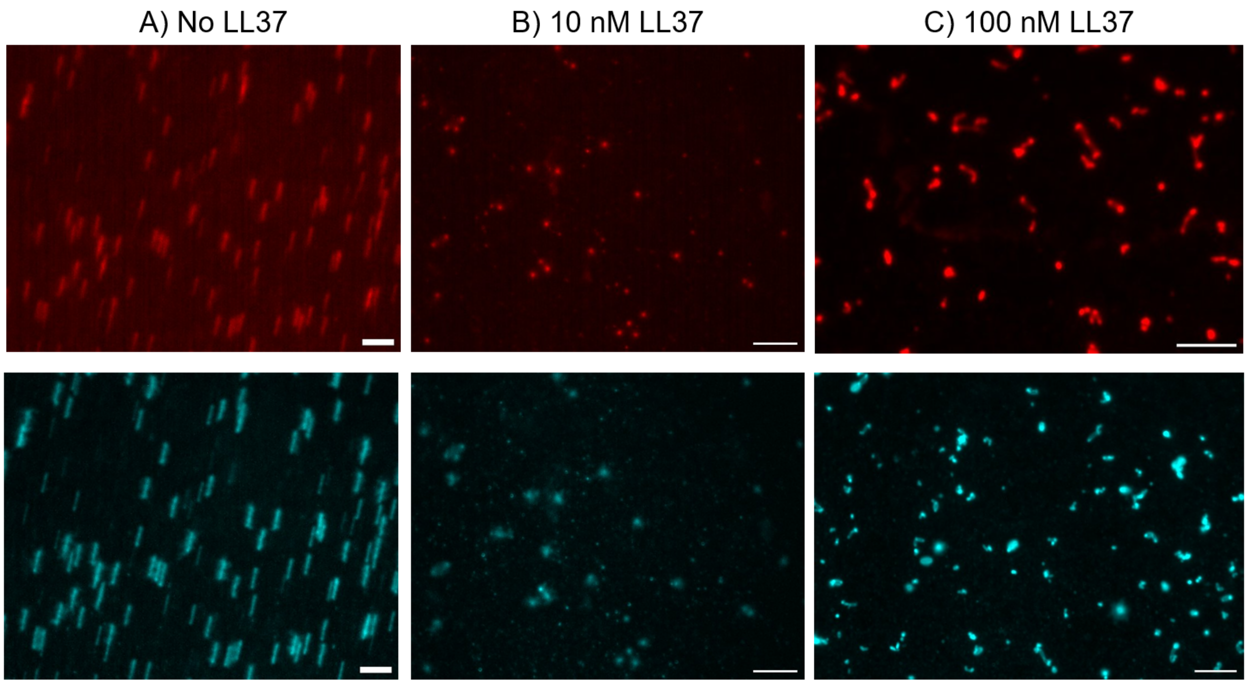


Figure S5: Pre-formed LL37-λDNA complexes form condensed structures. A) λDNA stained with 40 nM Sytox Orange was added to a flow channel and imaged in the absence of flow. DNA molecules were freely diffusing in solution as observed by grouped average intensity (top panel, green) and grouped standard deviation of a movie. B) λDNA pre-mixed with 10 nM LL37 showed freely diffusing behavior, with some DNA attached to the surface. C) λDNA pre-mixed with 100 nM LL37 showed multiple attachment points and condensed regions as observed the grouped average intensity (top panel). The grouped standard deviation (bottom panel) shows the mobility of regions between the condensed complexes. Scale bars are 5 µm.

Movie S1: Pre-formed LL37-λDNA complexes form condensed structures. No LL-37 added. (Pre-mix DNA with no_LL-37_crop1.avi)

Movie S2: Pre-formed LL37-λDNA complexes form condensed structures. 10 mM LL-37 added. (Premix DNA with 10 nM LL-37.avi)

Movie S3: Pre-formed LL37-λDNA complexes form condensed structures. 100 mM LL-37 added. (Premix DNA 100 nM LL-37.avi)

Movie S4: DNA curtains with 1 uM scLL-37 do not show any condensation (DNA Curtain 1uM scLL-37.avi)
